# A dynamic atlas of immunocyte migration from the gut

**DOI:** 10.1101/2022.11.16.516757

**Authors:** Silvia Galván-Peña, Yangyang Zhu, Bola S. Hanna, Diane Mathis, Christophe Benoist

**Author notes:** Address correspondence to: Christophe Benoist, Department of Immunology Harvard Medical School, 77 Avenue Louis Pasteur, Boston, MA 02115.

## Abstract

Dysbiosis in the gut microbiota impacts several systemic diseases. One possible mechanism is the migration of perturbed intestinal immunocytes to extra-intestinal tissues. Combining the Kaede photoconvertible mouse model and single-cell genomics, we generated a detailed map of migratory trajectories from the colon, at baseline and during intestinal and extra-intestinal inflammation. All colonic lineages emigrated from the colon in an S1P-dependent manner, dominated by B lymphocytes with a large continuous circulation of follicular B cells, which carried a gut-imprinted transcriptomic signature. T cell emigration was more selective, with distinct groups of RORγ^+^ or IEL-like CD160^+^ cells in the spleen. Gut inflammation curtailed emigration, except for DCs disseminating to lymph nodes. Colon emigrating cells distributed differentially to tumor, skin inflammation, or arthritic synovium, the former dominated by myeloid cells in a chemokine-dependent manner. These results thus reveal specific cellular trails originating in the gut, influenced by microbiota, which can shape peripheral immunity.

## INTRODUCTION

The gut is a complex organ, harbouring most of the organism’s microbiota, a constant flux of diet antigens, and a large number of immunocytes. Gut immunocytes face the unique challenge of maintaining tolerance to food and commensal antigens without compromising effective responses against pathogens. In addition, there is evidence that what happens in the intestines can influence systemic immune responses ^1,2^. The onset and progression of some extra-intestinal inflammatory diseases appear to be influenced by the intestinal microbiota ^3^. For instance, gut-colonizing Segmented Filamentous Bacteria (SFB) induce IL17-producing T cells that exacerbate disease in autoimmune mouse models of arthritis or multiple sclerosis ^4,5^. Several studies have reported perturbations in the intestinal microbiota of rheumatoid arthritis (RA), multiple sclerosis and type-1 diabetes patients ^6,7^, and the composition of the gut microbiome affects the responsiveness to immunotherapy in cancer patients ^8,9^. In addition, inflammation at distal sites such as the skin, the joints and the eyes can ensue in up to 50% of inflammatory bowel disease (IBD) cases ^10^.

The underlying mechanisms by which the gut is mediating this inter-organ communication remain incompletely understood. Several studies have demonstrated a role for metabolites and other bacteria-derived products ^11^, others have identified live bacteria as possible conduits ^12,13^. Another way in which the intestines might exert systemic influences is via migration of gut immunocytes. While we have some appreciation for cell migratory mechanisms and dynamics from lymphoid organs to the gut ^6^, our understanding of migration from the gut to lymphoid and other systemic organs is scanter. There is evidence that trafficking of several immunocyte populations from the intestines to the spleen and lymph nodes (LNs) takes place under steady-state ^14-17^. Intestinal microbiota-activated Th17 cells have a pathogenic role in multiple tissues in autoimmune mouse models of arthritis, uveitis, renal disease and encephalomyelitis ^4,5,18,19^. In addition, IgA^+^ plasma cells and FoxP3^+^ T regulatory cells (Tregs) of gut origin have protective roles in the brain and the pancreas ^20-23^, while intestinal innate lymphoid cells (ILCs) mediate anti-helminth defense and tissue repair in the lungs ^24^. If we seek ultimately to therapeutically target systemic effects of the microbiota in the context of disease, we must first build a solid understanding of these systemic migration pathways at baseline as well as in inflammatory conditions.

Kaede transgenic mice express a fluorescent protein that converts from green to red upon exposure to violet light, making them an ideal tool to *in vivo* track cellular migration minimally invasively ^25^. They have been previously used to temporally trace the migration of intestinal cells following photoconversion of the small intestine or ascending colon via surgical exposure ^19,26-29^, or in combination with a less invasive endoscopic system ^14,16^. While it was shown that immunocytes can exit the intestines, information on their destination and function has primarily concerned lymphoid tissues (mostly local LNs), or specific cell types in autoimmune models ^14,15,18,19,30,31^. Here we used a systems approach to obtain a global picture of the contribution of colon-derived immunocytes to circulation at homeostasis, and how this map is modified by inflammation or tumors in the gut or in destination organs. This work revealed that immunocytes emigrating from the intestines carried along a phenotypic memory of their origin and represented specific subsets in their destination organs, with cell-specific preferences for different inflammatory sites or tumors.

## RESULTS

### Gut immunocytes have high mobility and turnover

Colonoscopic illumination with violet light temporarily labels colon cells in photoconvertible Kaede mice in a short time window ^14-16,27^. To better understand the turnover of intestinal immunocytes, we tracked the proportion of photoconverted cells remaining in the descending colon at various times after the light pulse (Fig. 1A, B, gating in S1). This turnover reflected egress from the colon, but can also encompass cell death and proliferation. Tagged mononuclear phagocytes (MNPs), as well as T cells, disappeared in a bimodal fashion, a fastest drop happening in the initial 12-24hr, followed by a phase of slower decay, with 20-40% of the photoconverted populations still present after 3 days (Fig. 1A, B). In contrast, photo-tagged B cells declined faster, fitting a monophasic exponential decay, with very few red-labelled cells remaining after three days.

**Figure 1.**
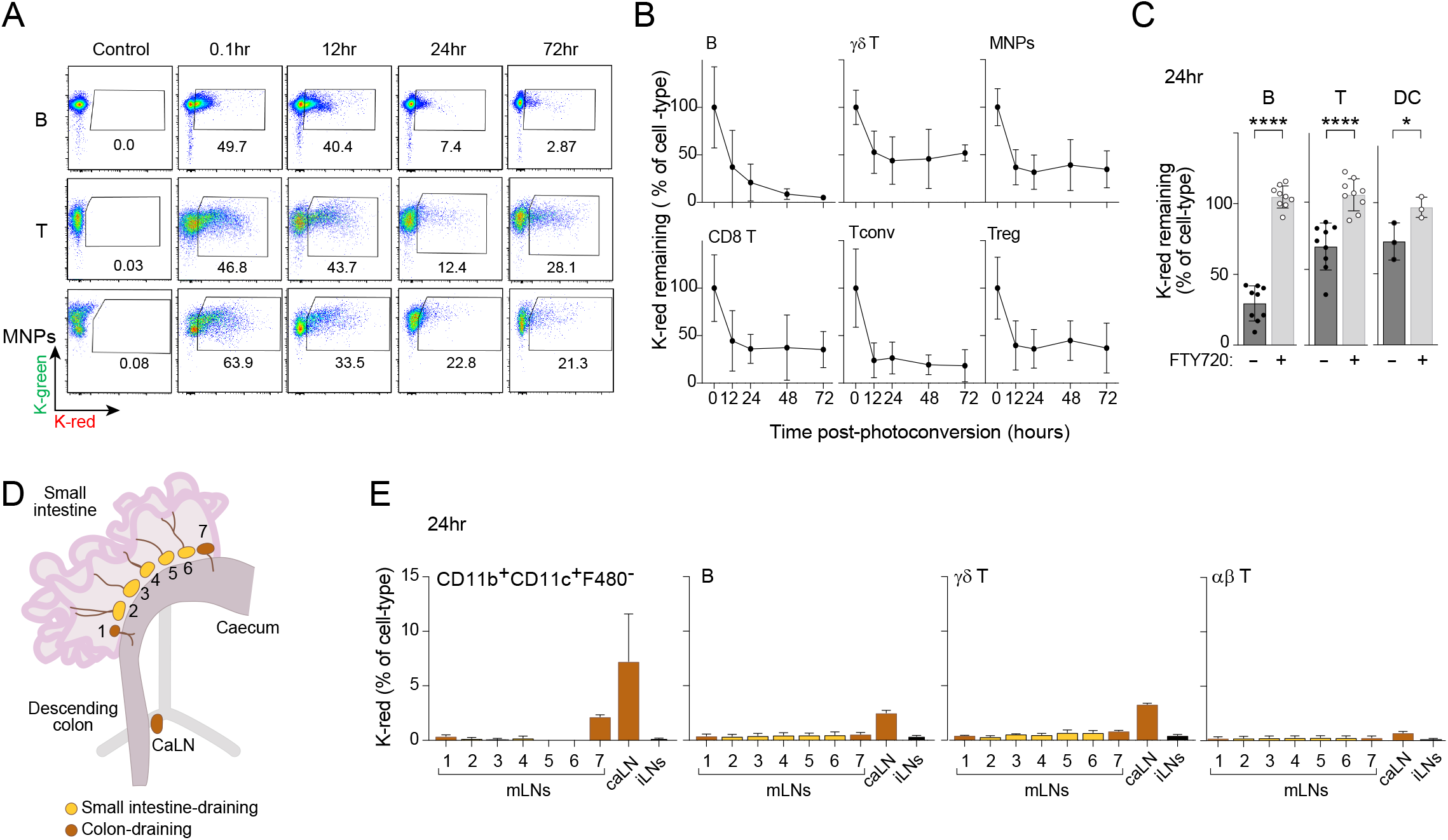
Turnover of colon immunocytes. A) Representative flow cytometry plots of photoconversion proportions of B, T and MNPs in the descending colon in unconverted (control) mice versus 10min, 12hr, 24hr, 72hr post colonic photoconversion. B) Egress rates of cells photo-tagged in the descending colon, measured by flow cytometry (n=4-6). Normalized to the average of time 0hr. C) Proportions of photo-tagged B, αβ T and CD11c^+^ DCs left in the colon 24hr post-photoconversion, with and without pre-treatment with FTY720 (1mg/kg). Normalized to the average of time 0hr. D) Schematic of gut draining LNs. E) Percentage of migratory Kaede-red populations across individual mLNs, caLN and iLN, 24hr post-photoconversion of the descending colon, measured by flow cytometry (n=4-6). All results from two to three independent experiments; Error bars indicate mean ± SD; *p < 0.05, ****p < 0.0001, unpaired t test.

We asked what mechanism underlies this fast turnover. Egress of T cells from the colon is S1P-dependent ^15,19^. Pre-treatment of Kaede mice with the S1PR functional antagonist FTY720 prior to photoconversion of the descending colon completely inhibited immunocyte turnover measured 24hr later, not just for T cells but also CD11c^+^ dendritic cells (DCs) and B cells (Fig. 1C). This result indicates that turnover of photo-tagged cells resulted from egress (as opposed to cell death or division) and that S1P was the main mechanistic controller.

Given this extensive egress from the colon, we first investigated local migratory paths. Mesenteric and caudal LNs (mLNs, caLN) are the site of direct lymphatic drainage for intestinal tissue, and classically the main location for antigen presentation to T cells by DCs ferrying microbial and food antigens from the gut ^32^. Drainage of the gut is compartmentalized, the distal colon being drained predominantly by the caLN and by a minority of the mLNs ^26,33,34^ (Fig. 1D). Examining individual nodes in the mLN chain one day after photoconversion of the colon in Kaede mice revealed a strong accumulation of MNPs of recent colon origin, CD11b^+^ CD11c^+^ dendritic cells (DCs) specifically, in the caLN (Fig. 1E). Immigrant DCs were also present in the most distal members of the mLN chain (#7 and #1), albeit to a lesser extent, but not in other mLNs, in agreement with prior studies. Photo-tagged B and T lymphocytes of colon origin followed a similar pattern of accumulation. It is worth stressing the high percentage of photo-tagged DCs in the caLN (10% of the DCs present in the caLN were in the colon the day before), much higher than in the spleen or subcutaneous LNs (inguinal (iLNs)), in line with the classic migration of DCs to draining LNs.

### Migratory immunocytes connect the colon with system-wide locations at steady-state

We examined the destinations of this migration, in lymphoid and non-lymphoid tissues. Indeed, 24 hr after photoconversion in the colon, we could detect photo-tagged red CD45^+^ cells in almost all tissues examined (obviously at much lower frequencies than in the colon, since colon-derived cells were diluted by local tissue-resident cells). The two notable exceptions were the thymus and the skin, where no migrant cells were detected (Fig. 2A). The latter supports the concept that the gut and skin immune systems are compartmentalized under normal conditions, even though both are microbiota-carrying barrier tissues ^35^. In the case of the thymus, which has been linked to the intestines via migratory DCs ^16,17^, colon-derived population may be diluted in large numbers of thymocytes, and thus below our level of detection. On the other hand, migration of colon-derived immunocytes to the lungs, liver and kidneys was surprisingly comparable with migration to lymphoid organs in terms of ultimate seeding frequency.

**Figure 2.**
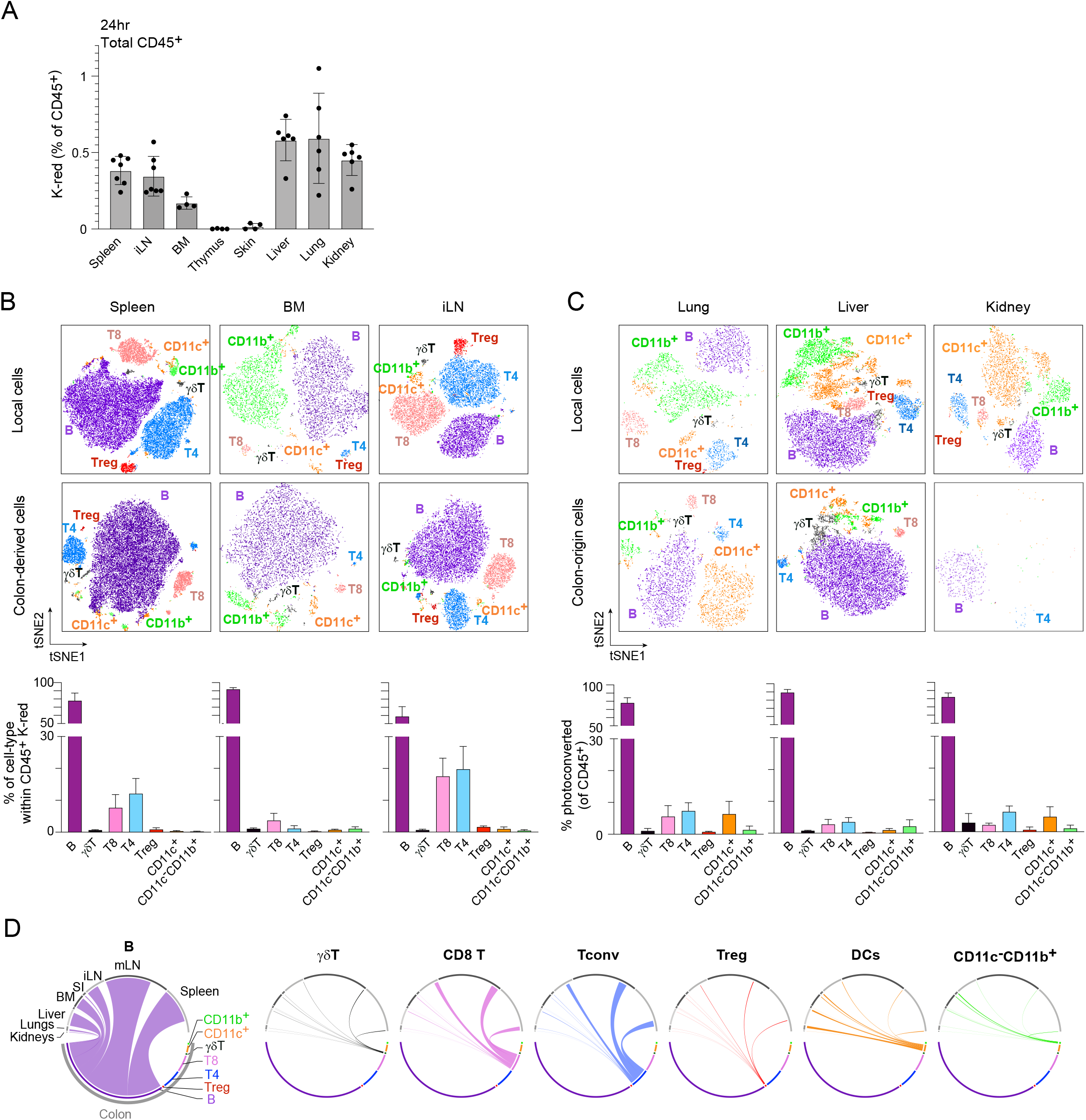
Immunocytes from the colon migrate into both lymphoid and non-lymphoid tissues. A) Percentage of CD45^+^ migratory Kaede-red cells across tissues, measured by flow cytometry 24hr following colon photoconversion. B and C) Local (Kaede-green) immunologic populations compared to colon-origin (Kaede-red) cells in (B) BM, spleen, iLN, (C) lung, liver and kidneys post colonic photoconversion by high dimensional flow cytometry. tSNE of CD45^+^ cells built on CD11c, CD11b, F4/80, γδTCR, αβTCR, CD8, CD4, CD25, CD19 (n=6). D) Chord diagrams depicting the numeric distribution of each Kaede-red population across all tissues examined (n=6). All results from three independent experiments; Error bars indicate mean ± SD.

To characterize the incoming colon-derived populations and compare them with their local tissue counterparts, we used high-parameter flow cytometry represented on a 2D tSNE projection (Fig. 2B, quantitated as a proportion of photoconverted CD45^+^ cells, bottom panels). In the spleen and bone marrow, B cells were the most dominant immigrant population. This dominance was less extreme in the LNs, where colon-derived T cells were better represented. When represented as a fraction of whole lineages, migrated cells represented between 0.2 and 0.5% of the total populations (Fig. S2), indicating that colon-derived cells contribute to every lineage in lymphoid organs. As the exception, although they represent a major fraction of the bone marrow cells, very few CD11c^-^CD11b^+^ cells were colon-derived, which is expected if one considers the bone marrow to be a locale of myeloid differentiation that should be somewhat shielded from environmental influences.

In non-immunological organs, B cells were again the dominant CD45^+^ immigrant population, much as in lymphoid organs (Fig. 2C). T cells were also present, with colon-derived Tregs making a larger contribution to the local niche than in the spleen (Fig. S2). Independently of proportional representation, compiling the tissues in which photo-tagged immunocytes that emigrate from the gut were found, revealed subtle but reproducible tissue preferences. MNPs in general migrated in greater numbers to non-immunological tissues than to LNs and spleen, in contrast to lymphocytes which did the opposite (Fig. 2D). Within lymphocytes, γδT cells and Tregs more closely resembled MNPs in their migratory preferences. Thus, our data showed widespread migration of immunocytes from the colon to both lymphoid and non-lymphoid tissues, mapping routes of inter-tissue connectivity. It may be worth emphasizing that the fractions observed result from a single pulse-label 24 hr earlier, and likely under-represent the cumulative numbers of colon-derived immunocytes in these tissues.

### B cells of colon origin bring a distinct transcriptome in the spleen

This widespread migration of gut immunocytes raised several questions relevant to their impact on systemic immunologic function: how do these gut migrants differ from their counterparts in destination tissues? Do they express gut-imprinted phenotypic and functional programs? To compare colon-derived vs local populations in a broad and unbiased manner, we performed single-cell RNA sequencing (scRNAseq) on total CD45^+^ splenocytes of Kaede mice at different time points after colonoscopic photoconversion (Fig. 3A; sorting equivalent numbers of Kaede-green and red cells, every timepoint from 2 individual mice, 13,833 cells altogether). The use of DNA-coded antibody hashtags ^36^ enabled us to profile twelve samples in the same run, providing an ideal comparison between immigrant and resident cells. Dimensionality reduction and projection on a Uniform Manifold Approximation and Projection (UMAP) revealed all the immunologic lineages expected in the spleen (Fig. 3B), identifiable with the usual marker genes (Fig. S3A) and represented across both resident and migrant populations (Fig. 3B; Fig. S3B).

**Figure 3.**
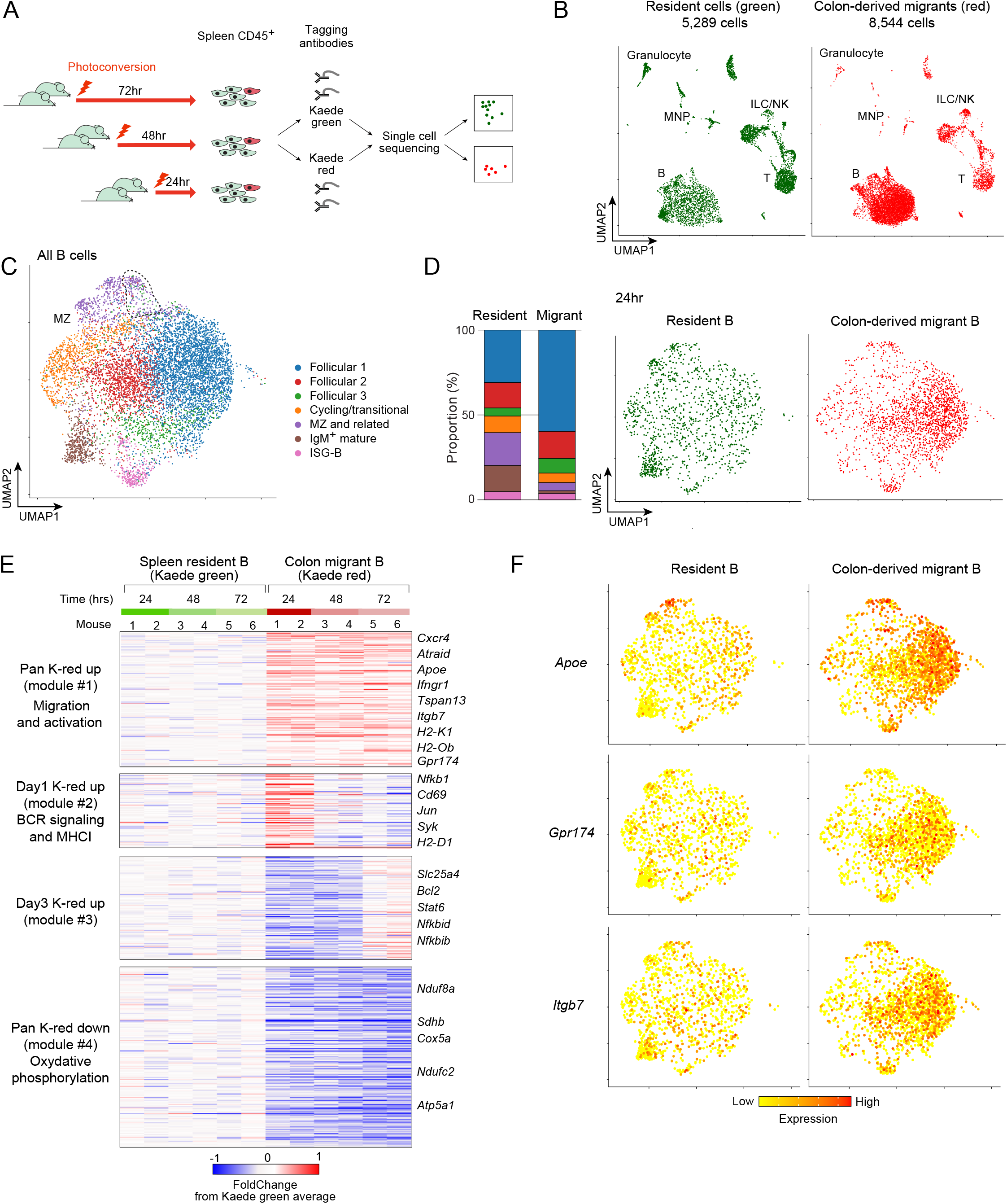
Transcriptome of colon-derived B cells. A) Single cell experiment schematic. Splenic cells were isolated from mice photoconverted at various points and sorted as CD45^+^ Kaede-green^+^ or CD45^+^ Kaede-red^+^, labelled with hashtag antibodies and sequenced in a single lane. Sample demultiplexing was performed computationally. B) scRNAseq analysis of total CD45^+^ cells in spleen. UMAP representation, color-coded by cell identity. C) UMAP representation of B cells selected from (B). Color coded by *k*-nearest neighbour (knn) cell clusters (total 8440 cells). D) Distribution of 24hr post-photoconversion Kaede-green vs Kaede-red cells across the UMAP in (C) (1194 Kaede-green cells, 2002 Kaede-red cells). E) Clustered heatmap of differentially expressed genes between Kaede-red and Kaede-green cells within the top four most migrant-enriched clusters from (D), across all three timepoints (355 genes, *p <* 0.01). F) Gene expression of selected markers across Kaede-green and Kaede-red cells (same plots as in D).

In agreement with our flow cytometry data, B cells constituted a large portion of the migratory compartment. To examine more finely the phenotypic state of these migratory cells, we first re-clustered them on their own, Louvain clustering distinguishing seven different B-cell types (Fig. 3C), that coincided with specific B cell markers (Fig. S3C, S3D). These subsets included distinctive marginal zone (MZ), mature IgM^hi^, a small population with high expression of interferon signature genes (ISG) (hereafter called ‘ISG-B’, by analogy with comparable T cells ^37-39^), and a large contingent of naive follicular B cells. Newly arrived colon-derived B cells (24hr after photo-tagging) were markedly missing from the IgM^hi^ mature group, and distinct from the main MZ subset (Fig. 3D). Migrants mostly belonged to the follicular clusters, with a dominance of IgD^+^CD23^+^CD21^lo^ B follicular-like cells. A differential density plot between the colon-derived and resident cells also reflected these differences, which remained largely unchanged at a population level for 72hr (Fig. S3E), although the distribution within the follicular clusters did evolve somewhat. These particularities were confirmed by flow cytometry (Fig. S3F), and in an independent scRNAseq experiment (Fig. S3G). Colon-derived B cells also included a surfeit of ISG-B cells, of interest considering recent reports highlighting the role of microbiota-induced type-I IFN ^40,41^.

To identify gut-imprinted gene-expression programs we performed differentially expressed genes (DEG) analysis between the migratory (red) and local (green) splenic B cells. To avoid differences merely stemming from variable cluster representation, DEGs were determined within each of the four most represented clusters, selecting genes with significantly different expression between migrant and resident B cells in every cluster (at p < 0.01). The resulting group of 355 DEGs distinguished colon-derived B cells (Fig. 3E). Some remained differential at all timepoints (module #1 in Fig. 3E), including the genes encoding the major gut-homing integrin Itgb7, and perhaps less expectedly transcripts encoding the chylomicron apolipoprotein ApoE (Fig. 3F) or the lysophosphatidylserine receptor Gpr174, whose functions in B cells are less obvious ^42^. Expression of some transcripts normalized after a day or two (module #2), while other gene-sets remained under-expressed in immigrant cells (modules #3 and #4). Together, these results revealed a circulatory population of gut-derived B cells with gut-imprinted characteristics that were still distinguishable three days after exit from the colon.

### Multiple populations of colon-derived T cells are present outside the gut

Migratory T cells told a somewhat different story from that of B cells. Re-clustering the scRNAseq data to focus on T cells (Fig. 4A, Fig. S4A-B) resolved the expected populations of CD4^+^ and CD8^+^ T cells (naïve and effector) as well as more distinctive subsets (FoxP3^+^ Treg, ISG-T ^37-39^). Many of the colon-derived cells corresponded to the main naïve CD4^+^ and CD8^+^ T cell groups, however the comparison of Kaede-green and red T cells revealed several activated/memory populations that clearly differentiated spleen-resident and colon-derived cells (arrows in Fig. 4B, detailed in Fig. 4A,C)). iNKT cells, identified by their invariant TRAV11-TRAJ18 TCRα chains and by a distinct set of transcripts (Fig. S4C), were essentially absent from migrant cells (in keeping with the observation that iNKTs are quite rare in the colon lamina propia (LP)). In contrast, two distinct clusters were almost exclusively represented by recent immigrant Kaede-red cells. One of these (‘Rorγ T’ in Fig. 4A) expressed *Rorc* transcripts and included cells expressing either αβ or γδ TCRs (Fig. 4C). In addition to *Rorc*, these cells expressed *Cd44, Tmem176, Cxcr6, Il2ra* (but not *Foxp3*) and lacked *Cd27*, characteristics of Th17 cells (Fig. S4A-B). Although transcripts encoding IL17 were scarce, these characteristics pointed to colon-derived cells that could represent the conduits of the influence of Th17-inducing microbes on extra-intestinal immunity ^4,5,19^.

**Figure 4.**
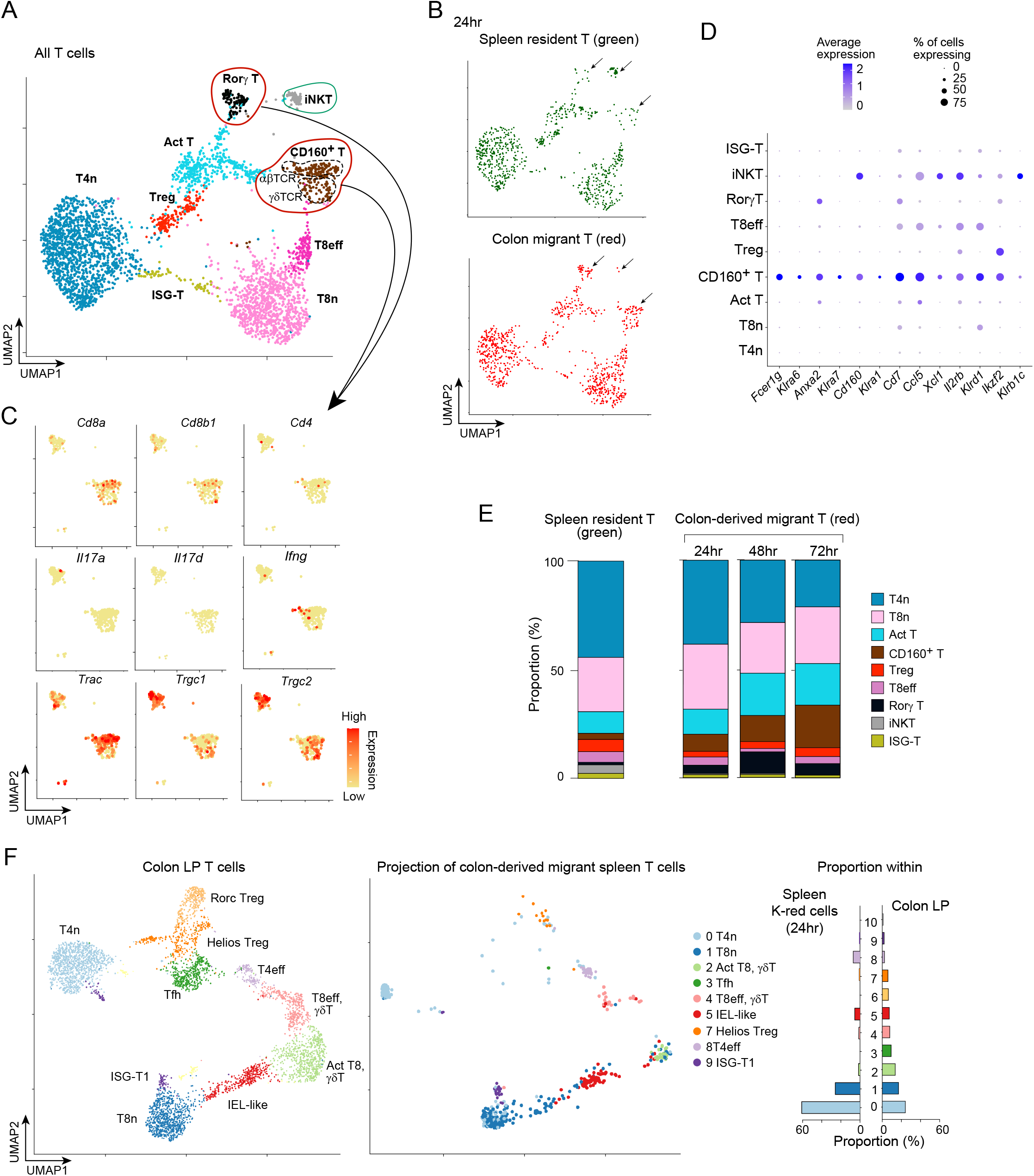
Distinct colon-derived migratory T cell populations. A) UMAP representation of T cells selected from 3B. Color-coded by knn cell clusters (total 3500 cells). B) Same plot as in (A), with Kaede-green vs Kaede-red distribution 24hr post-photoconversion (782 Kaede-green cells, 852 Kaede-red cells). C) Gene expression of selected markers across the ‘Rorγ T’ and ‘CD160^+^ T’ clusters from (A). D) Differential marker genes for the CD160^+^ T cluster. E) Percentage quantification of the distribution of the Kaede-green and Kaede-red cells across clusters per timepoint. F) scRNAseq analysis of total CD45^+^ cells from cecum. UMAP representation, color-coded by cell identity (n=2) (left); Kaede red T cells projected onto the dataset from (F) UMAP (right).

The other population dominated by cells of recent colonic origin (‘Cd160^+^ T’ in Fig. 4), was a *Cd160*-expressing cluster that also included both αβ and γδT cells. They expressed *Cd8a* more than *Cd8b1*, evoking CD8αα intra-epithelial lymphocytes (IEL) in the gut ^43^. The possibility of such a relationship was strengthened by the expression of several other markers typical of IELs such as *Fcer1g, Il2rb* and several *Klr* family members (but not *Klrb1c*, encodes NK1.1) (Fig. 4D, Fig. S4A). The expression of a broader IEL gene signature (from ^44^) was also biased for this population (Fig. S4D). Over time, while the main naïve CD4^+^ and CD8^+^ subsets faded, likely because they represented circulatory populations that happened to be in the colon at the time of photo-tagging, the IEL-like population as well as the other populations enriched in colon-derived cells remained stable in the spleen for multiple days (Fig. 4E).

We then asked the inverse question: could we identify, in a scRNAseq dataset of intestinal T cells, those that gave rise to migratory populations in the spleen. We generated a scRNAseq dataset of cecum LP T cells (Fig. 4F, left) and used a transfer-learning algorithm ^45^) to project onto these coordinates the Kaede-red immigrant cells from the spleen data. The cells that could be confidently assigned (Fig. S4E) mapped only to specific T cell subsets (Fig. 4F, middle and right panels). Other than the expected naïve CD4^+^ and CD8^+^ populations, and ISG-T cells, the majority of cells mapped to two clusters: effector CD4^+^ T cells (‘T4eff’) and an IEL-like effector population (‘IEL-like’), marked by expression of *Cd160, Eomes* and multiple Klr genes (*Klrc1, Klrb1c, Klrc2, Klra7, Klre1*) (Fig. 4D). Other populations, like T follicular helper (‘Tfh’) cells, were not represented in colon-derived cells in the spleen, despite being a sizeable subset in the colon. Thus, there is specific export of activated T cell subsets from the colon, some potentially related to Th17, but also other more unconventional subsets of γδ and CD8^+^ T cells.

### Intestinal perturbations modify the systemic flux of colon-derived populations

Having established in fine detail the landscape of cells migrating from the colon at baseline, we then investigated how perturbations in the gut affected this traffic. In mammalian organisms, the colon harbours the largest population and diversity of microbes, which led us to question their influence on migratory populations. We treated Kaede mice with a broad-spectrum antibiotic cocktail for 10 days (Fig. 5A), long enough to gravely reduce bacterial density (>10^4^-fold reduction in cultivatable bacteria in feces, both aerobic and anaerobic) yet not completely reshuffle the local immunological ecosystem (Fig. S5A). The migration to spleen and lymph nodes of immunocytes photo-tagged in the colon was significantly reduced in antibiotic-treated mice when compared with controls (Fig. 5B, no photoconversion differences S5A), significantly for B cells and γδT cells, but with clear trends for other lineages as well. Egress from the colon was slightly reduced, albeit not as significantly (Fig. 5C), thus indicating that migration from the colon was influenced by the microbiota.

**Figure 5.**
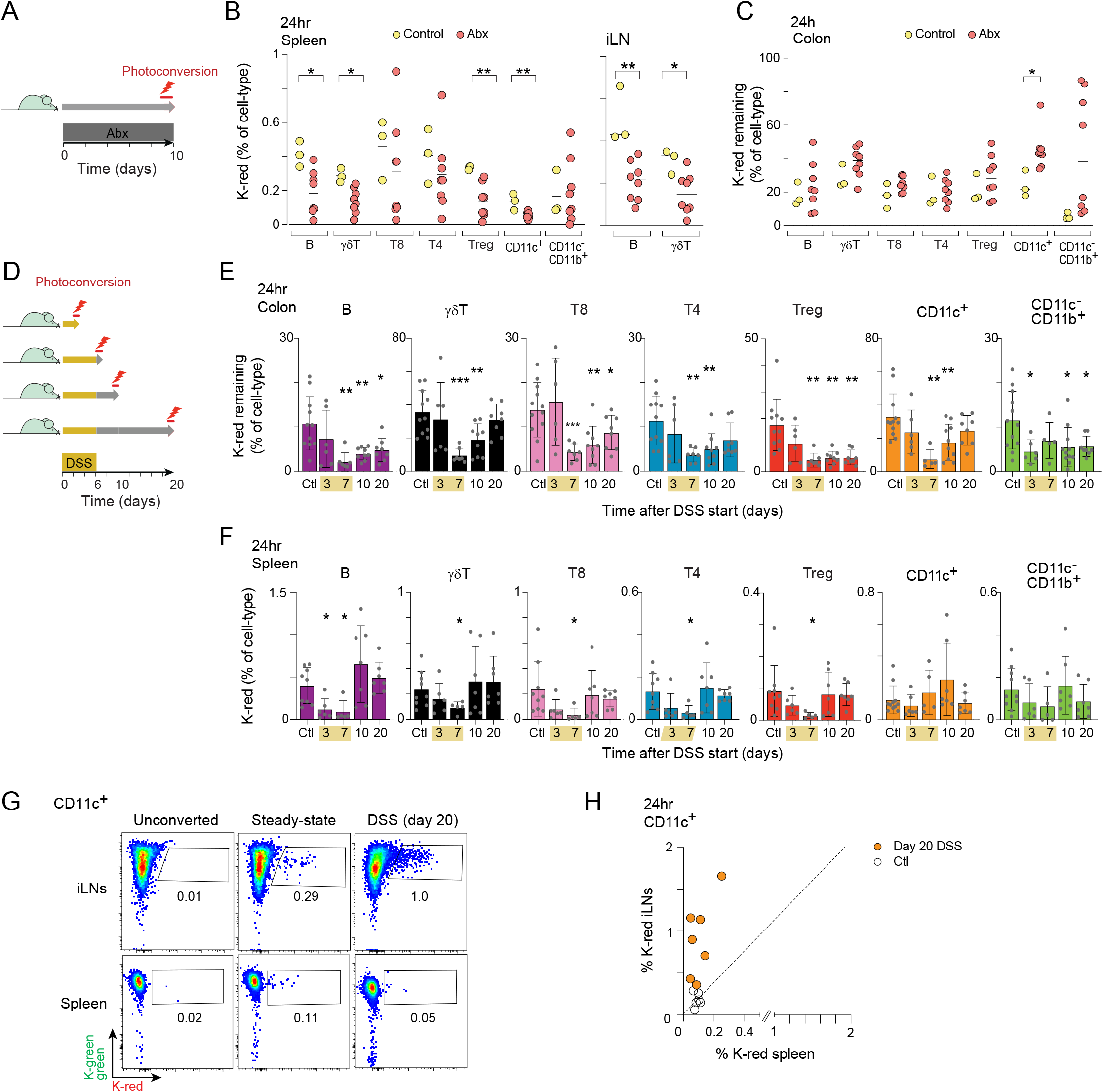
Intestinal perturbations significantly alter systemic migration. A) Schematic of the experimental set-up for antibiotic-treated mice. B) Percentage of colon-origin Kaede red populations in the spleen and iLN, 24hr post-photoconversion of the descending colon in mice treated with an antibiotic cocktail for 10 days (VGCA, vancomycin, gentamycin, clindamycin and ampicillin). C) Proportions of immunocytes left in the colon 24hr post-photoconversion of the descending colon in mice treated with an antibiotic cocktail for 10 days (VGCA). D) Schematic of the experimental set-up for DSS-treated mice. E) Proportions of immunocytes left in the colon 24hr post-photoconversion in control mice (Ctl) as well as different stages of DSS-induced colitis. F) Percentage of colon-origin Kaede-red populations in the spleen, 24hr post-photoconversion of the descending colon at different timepoints of DSS-induced colitis. G) Representative flow cytometry plots of the CD11c^+^ population in the spleen and iLN of control mice (steady-state) vs mice on day 20 post initial DSS administration, 24hr post-photoconversion of the descending colon. H) Correlation between Kaede-red CD11c^+^ in iLN (y-axis) and Kaede-red CD11c^+^ in spleen (x-axis) in control vs mice on day 20 post initial DSS administration, 24hr post-photoconversion of the descending colon. All results from two to four independent experiments. Each dot represents a mouse, mean is marked. *p<0.05; **p<0.01, ***p<0.001, unpaired t-test.

Reciprocally, we asked what happens to traffic during intestinal inflammation, a question related to extra-intestinal manifestations of IBD. We used the dextran sodium sulphate (DSS)-induced model of colitis, and photoconverted the distal colon at various stages of disease, from very early (48hr after initiation of DSS) to well into the resolution phase (2 weeks after cessation of treatment; Fig. 5D). As illustrated in Fig. 5E, the turnover of photo-tagged cells from the colon (assessed as above, 24hr after photoconversion) was greatly accelerated during full-blown inflammation (day 7), mostly returning to baseline values after recovery (day 20). This faster turnover was not a reflection of photoconversion efficiency (Fig. S5B). It was also not reflected as increased systemic migration, as measured in the spleen - indeed the opposite. With full-blown colitis (day 7), there was a 2-to 8-fold drop in the proportions of recent immigrants from the colon across all lineages (Fig 5F). Here again, migration returned to mostly normal levels after cessation of DSS (day 20). Thus, the inflammation induced by DSS led to a faster turnover of local immunocytes, likely by cell death and/or proliferation but perhaps also by shedding ^46^, and to a decreased seeding of extra-intestinal tissues.

Unexpectedly, we observed that migration of DCs to the iLNs was markedly increased, constituting up to 1% of the total DC pool in the lymph node on day 20 (Fig. 5G). This biased migration was not observed in the spleen where the proportion of immigrant DCs remained similar to control mice (Fig. 5F,H). No other populations showed such a bias, with perhaps the exception of γδ T cells, although to a far lesser extent (Fig. S5C). Thus, perturbations of the intestinal milieu significantly rewired cell transport.

### Immunocyte migration from the colon to systemic sites of inflammation

Finally, we asked the reverse question: how does inflammation in the destination organs affect the patterns of migration? We addressed this question in disease models that, while not directly affecting the gut, have previously been reported to have a connection to the gut microbiota: K/BxN arthritis ^4,6,47^, the MC38 tumor ^48-50^ and delayed-type hypersensitivity (DTH) to keyhole limpet hemocyanin (KLH) in the ear-skin ^51^ (Fig. 6A). Once inflammation or subcutaneous tumors were established, cells in the distal colon were photo-tagged by colonoscopy as above and migration to inflamed sites was quantitated 24hr later. Colon egress, as measured by percentage of remaining Kaede-red immunocytes, was unaffected in any of the models, except for an inconstant trend towards reduced MNP egress in arthritic mice (Fig 6B, Fig. S6A). Similarly, immunocyte migration from the colon to the spleen appeared largely unaffected in the disease models (Fig 6C, Fig. S6B).

**Figure 6.**
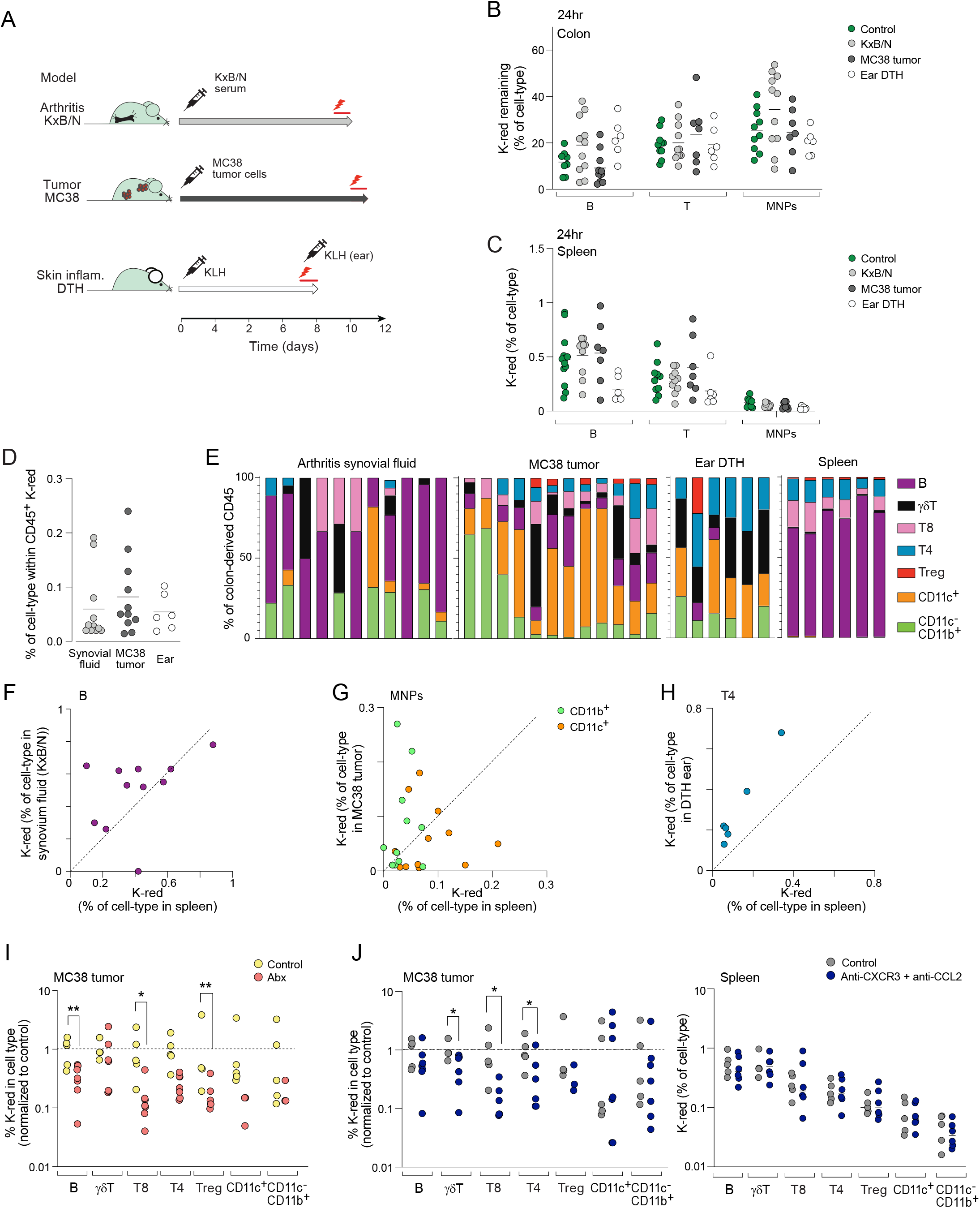
Immunocyte migration from the colon to distal diseased sites. A) Schematic of the experimental set-up for the KxB/N serum-transfer arthritis model, MC38 subcutaneous tumor model and the skin hypersensitivity DTH model. B) Proportions of immunocytes left in the colon 24hr post-photoconversion in control mice and mouse models of arthritis (KxB/N, day 10), MC38 tumors (day 11), and DTH (day 8). C) Percentage of colon-derived Kaede-red populations in the spleen, 24hr post-photoconversion of the descending colon in the same mice as in (B). D) Percentage of CD45^+^ cells 24hr post-photoconversion of the descending colon in the synovial fluid of KxB/N mice, MC38 tumors and the inflamed ear of DTH mice. E) Identity of the colon-derived immunocytes, as measured by flow cytometry 24hr post-photoconversion of the descending colon in synovial fluid of KxB/N mice, MC38 tumors and the inflamed ear of DTH mice. Each column represents a mouse. F) Correlation between Kaede-red CD19^+^ B cells in the synovial fluid of KxB/N mice (y axis) and the spleen from the same mice (x axis), 24hr post-photoconversion of the descending colon. G) Correlation between Kaede-red MNPs in MC38 tumors (y axis) and the spleen from the same mice (x axis), 24hr post-photoconversion of the descending colon. H) Correlation between Kaede-red Tconv (CD4^+^ CD25^-^) in the inflamed ear of DTH mice (y axis) and the spleen from the same mice (x axis), 24hr post-photoconversion of the descending colon. I) Percentage of colon-derived Kaede-red cells in MC38 tumors of antibiotic-treated mice (11 days, VGCA), 24hr post-photoconversion of the descending colon. Data normalized to the average of each control. J) Percentage of colon-derived Kaede-red cells in MC38 tumors of mice pre-treated with anti-CXCR3 (250μg/mouse) and anti-CCL2 (250μg/mouse) antibodies, 24hr post-photoconversion of the descending colon. Data normalized to the average of each control. All results from two to four independent experiments. Each dot represents a mouse, mean is marked. *p<0.05; **p<0.01, unpaired t-test.

Thus, inflammation in extra-intestinal tissues did not markedly resize exiting from the colon. On the other hand, migration to the sites of inflammation showed distinct characteristics. The overall contribution of recent emigrants from the colon was roughly equivalent in all three affected tissues (Fig. 6D), but the identity of the cells infiltrating each tissue was quite different. In in the synovial fluid, colon-derived Kaede-red cells were mostly composed of B cells, in percentages generally comparable with those for the spleen (Fig. 6E, F). In contrast, MC38 tumors were predominantly infiltrated by myeloid cells from the colon. CD11c^-^CD11b^+^F480^+^ macrophages were the dominant colon-derived population (Fig. 6E), which was intriguing because macrophages were not a dominant migratory population elsewhere. Indeed, comparison of immigrant macrophage frequencies in the tumor and spleen of the same mice showed a clear bias towards the tumor (Fig. 6G). Given the influence of macrophage subtypes on tumor progression, this influx suggests a possible root for the influence of the microbiome on tumor immunotherapy ^8,9^, and that tumor-infiltrating macrophages can have more diverse origins than simply blood monocytes ^52^ Finally, the inflamed DTH ear recruited a mix of immunocytes, but was the site with the relatively largest component of CD4^+^ Tconv and γδT cells (Fig. 6E, H). This homing contrasts with the absence of migration to the skin at baseline, indicating that inflammation carves new destinations for colon-derived immunocytes.

We had previously observed that steady-state systemic migration of these gut-imprinted populations could be reduced by diminishing bacterial load via antibiotics (Fig. 5B). Mice treated with a broad-spectrum antibiotic cocktail for the duration of tumor growth also showed reduced infiltration of colon-derived immunocytes into the tumors, with γδ T cells being one notable exception (Fig. 6I). Thus, by targeting the intestinal commensal bacterial load, we could manipulate entry of colon-derived populations at a distally perturbed site. We then questioned the reverse: would targeting known mechanisms of tumor entry also affect colon-derived populations? Treating the mice at the time of photoconversion with blocking antibodies against CXCR3 and CCL2, both reportedly used by immunologic populations to infiltrate MC38 tumors ^53^, reduced entry of colon-derived T cell populations into the tumors, but had no effect on proportions in the spleen (Fig. 6J). MNPs were not affected however, perhaps as a result of redundancy in their chemokine receptors. These data, together with the S1PR-dependent colon egress, would indicate that once these immunocytes exit the intestines, they at least in part use the same mechanisms as other circulatory populations to traffic to and enter distal tissues.

## DISCUSSION

Half a century ago, Hall and Smith postulated that any immune response must incorporate cells from the intestines, arguably the largest immunologic organ ^54^. While cell trafficking between lymphoid organs has been studied for decades ^55,56^, it is only more recently that several studies have shown that cell migration from the intestines does occur ^14,17,20,24,28^. We set out to map in detail these migratory paths from the gut to other organismal locations, their cellular and molecular specificity, and how these patterns vary under immunologic challenges. We uncovered a constant conveyor belt of immunocytes fanning out from the colon to extra-intestinal sites, one with clear cell-specificity in terms of the emigrating subsets and their preferential destinations, ferrying antigen-loaded cells and specific immunocyte phenotypes including expression of receptors characteristic of gut-experienced cells.

Numerically, B cells dominated the traffic from the colon, and were found at every destination. The majority of these immigrant B cells were of the common follicular type, and it is not obvious what these cells might bring to the target tissues (although they did carry a genomic signature of their gut residence). It might be that B cell emigration represents merely the exit phase of a strong flux of large numbers of B cells through the gut in order to select and exploit microbe-specific immunoglobulins in the B cell repertoire (or an evolutionary remnant of the bursa?). In contrast, migrant T cells included more specifically differentiated cells, carrying the imprint of microbial influence. DCs moving from the colon to the draining caLN stood out in terms of proportional representation (10% of the caLN DC pool came from the colon over a 24 hr period, dwarfing any other frequencies; incidentally, the results of Fig. 1E highlight the importance of carefully splitting mLNs when analysing intestinal lymphatic drainage, to avoid dilution by irrelevant locales). This corresponds to the textbook ferrying of antigen by DCs to draining LNs, but it is interesting that DCs also trafficked to other LNs, albeit in much smaller numbers, especially in the context of gut inflammation.

The combined single-cell profiling, with over-sampling of the Kaede-red populations, brought forth distinct subsets that would otherwise have been missed, being aggregated with other Teff cells by the clustering process. One such population of gut origin was the CD160^+^ T cell group, which included both αβT and γδT cells. Their gene expression resembled that of IEL populations (e.g. *Cd8a, Itgb7*, Klr genes), which is intriguing given that IELs are generally considered to be tissue-resident. On the other hand, a recent study documented the emigration of several tissue-resident, non-classical T cells to local draining lymph nodes ^30^. Thus, while the majority of IELs may be tissue-resident, some may enter the general circulation. As for their function, one might speculate that their unique position within the epithelium, unique access to the intestinal antigenic load, and their innate-like responsiveness might make them potent regulators outside of the gut ^57^.

Another distinct population of colon origin uncovered in the spleen was the RORγ^+^ T cell, also composed of both αβ and γδ T cells. IL-17-producing T cells in the gut, which depend on the microbiota ^58-61^, help maintain local homeostasis ^62-64^. However, they also exhibit pathogenic roles in extra-intestinal autoimmune disorders ^4,5,18,19,31^, with previously reported migration of gut Th17 in these pathological settings. It has been proposed that gut Th17 cells fall into two classes: homeostatic and pathogenic ^65^. But caution should be exercised in using dichotomic models which may oversimplify more continuous gradients ^37^. Uncovering a robust emigration of RORγ^+^ T cells from the gut in unchallenged mice is interesting in this regard. The pathogenic gut-brain or gut-joint axes may result from this baseline conveyor belt, but be corrupted by a perturbation at its gut origin. Alternatively, rather than an overall change in phenotype of these migratory cells, pathogenicity may stem from a change in the balance between populations of γδ vs αβ T cells.

Also worth discussing are the ISG-T and ISG-B phenotypes, previously described in T cells ^37-39^, but also seen here to exist in B cells: a small subset of otherwise unremarkable cells characterized by unusually high ISG expression (whether from a particular sensitivity to normal levels of IFN or from high exposure in a particular microenvironment). Our results suggest that ISG-T cells in systemic locations might actually derive from the gut, an interesting notion given that the mucosa is a frequent point of entry for pathogens. Might these cells with potentially microbe-specific receptors, but protected by high ISG levels, be advantageous in extra-intestinal locales? In addition, these observations raise the possibility of involvement in systemic pathogenesis, e.g. in B cell pathologies that involve IFN, like systemic lupus erythematosus ^66^.

Extra-intestinal manifestations of inflammatory bowel disease are a good example of cell migration as a possible mechanism by which gut dysbiosis becomes a systemic problem. Migration of T cells from the intestines to the liver during colitis has been implicated in mediating liver pathogenesis ^67^, and a recent report highlighted changes in the antigen-specificity of skin T cells following colitis ^29^. Interestingly, the skin was one of very few tissues we could not detect any gut-derived immunocytes, thus implying that pathogenesis cannot be driven solely by corruption of existing axes of cell transport, but that new ones can form as a result of gut dysbiosis, or of local inflammation (as in the DTH ear).

Our results show that gut-educated populations also homed to sites of inflammation, with a cell specificity that varied with the type of lesion. It was in DTH lesions that T cells were most present, perhaps not coincidentally since DTH is a T cell-mediated disease. The relative dominance of myeloid cells in MC38 tumors was intriguing. Tumor-infiltrating MNPs are thought to be largely of monocyte origin, but these results indicate a sizeable flow of MNPs from the gut, which may relay the documented effects of the gut microbiota on tumor sensitivity to checkpoint blockade ^8,9^. Thus, understanding the migratory dynamics of gut immunocytes in different disease states, as we have begun to do here, might provide new approaches to the modulation of immunologic disorders and their interplay with the gut and its microbiota.

## ACKNOWLEDGEMENTS

We thank Dr. A. Magnuson and Dr Ramanan for advice, B. Vijaykumar and D. Mallah for help with computational analyses, A. Ortiz-Lopez for help with experiments and K. Hattori for mice. This work was supported by NIH grants AI125603 and AR070334, the JPB Foundation, and in part by an SRA from Evelo Biosciences. SG-P was supported by a fellowship from the European Molecular Biology Organisation (ALTF 547-2019), BSH was partially supported by a Deutsche Forschungsgemeinschaft fellowship (HA 8510/1)

## MATERIALS AND METHODS

### Mice

Kaede transgenic mice were originally obtained from O.Kanagawa (RIKEN, Wako, Japan), and bred on a C57Bl/6 background in our specific-pathogen-free facility. All experimentation performed following the animal protocol guidelines of Harvard Medical School (protocol IS-2954; 0196-3)

### In vivo treatments and disease models

Mice were pre-treated via intraperitoneal (ip) injections with either S1P receptor antagonist FTY720 (1 mg/kg, Cayman Chemical), anti-CXCR3 antibody (250 μg/mouse, BioXcell), anti-CCL2 antibody (250 μg/mouse, BioXcell) or anti-armenian hamster IgG (Invitrogen), at the time of endoscopic photoconversion.

For antibiotics treatment, vancomycin (0.5 g/L, VWR Life Science), gentamycin (0.5 g/L, VWR Life Science), clindamycin (0.5 g/L, alfa aesar) and ampicillin (1 g/L, Sigma) were dissolved in drinking water with sweetener and given to the mice for 10 days. For DSS treatment, 2.5% DSS (Thermo Scientific) was provided in the drinking water for 6 days.

For tumor induction, MC38 cells (obtained from Dr Arlene Sharpe, Harvard Medical School) were first cultured *in vitro* in DMEM supplemented with 10% FCS and 1% L-glutamine for a week, prior to subcutaneous injection of anesthesized mice with 1 × 10^6^ cells in 100 μL PBS.

K/BxN serum-transferred arthritis was induced by ip injection of 150μL pooled serum from 8-week-old K/BxN mice on days 0 and 2. Arthritis development was assessed by visual inspection of the paws and caliper (Kafer, 10 mm range with flat anvils) measurement of ankle thickness.

For the DTH model, mice were anesthesized, backs shaved and injected with KLH (Sigma, 1 mg/mL) emulsified in complete Freund’s adjuvant (CFA) emulsion on four different back sites (50 μL altogether). 8 days later, the mice were injected with 10 μL of KLH (dissolved in sterile PBS, 1 mg/mL) into one of the ears intradermally.

### Photoconversion procedures

Colon was photoconverted as previously described ^14^. Briefly, mice were first anesthetized with ketamine:xylazine (10 mg/kg:2mg/kg i.p). A custom-built fiberoptic endoscope (ZIBRA Corporation) was coupled to a handheld 405 nm blue purple laser (≤ 5mW) via an in-house custom-made connection device (fixed mounts from ThorLabs). After cleansing the colon of fecal pellets with PBS, the fiberoptic endoscope was inserted through the anus to a depth of 3 cm. The laser was switched on, exposing the inner colon to violet light (3.5 mm beam diameter). Subsequently, the endoscope was gently retracted, pausing at 2 mm increments for 30 s light pulses at each interval (for a total of up to 10 min).

### Immune cell isolation from tissues

#### Spleen, bone marrow and lymph nodes

Immunocytes were released by mechanical disruption followed by filtering and washed in RPMI containing 5% fetal calf serum (FCS). Red blood cells in the spleen and bone marrow were lysed prior to filtering using ACK lysing buffer (Gibco, ref A10492-01).

#### Colon and small intestine

Immunocytes were isolated as previously described ^28^. Briefly, intestines were cleaned (Peyer’s patches removed in the case of the small intestine), and treated with RPMI containing 1 mM DTT, 20 mM EDTA and 2% FCS at 37°C for 15 min to remove epithelial cells. They were then minced and dissociated in collagenase solution (1.5 mg/mL collagenase II (Gibco), 0.5mg/mL dispase (Gibco) and 1% FCS in RPMI) with constant stirring at 37°C for 40 min. Single cell suspensions were filtered and washed with RPMI containing 5% FCS.

#### Liver, lungs and kidneys

mice were first perfused with 5mL of ice-cold PBS through the heart’s left ventricle. Tissues were minced and dissociated in collagenase solution (0.5 mg/mL collagenase IV (Gibco), 150μg/mL DNase I (Sigma) and 1% FCS in DMEM) and incubated in a water bath at 37°C with constant shaking for 40 min. Digested tissues were filtered and washed in 2% FCS. For liver and kidneys, immunocytes were enriched by Percoll (GE Healthcare) density centrifugation (36%, 800 x g for 10 min). For lungs, red blood cells were lysed using ACK lysis buffer.

#### Thymus

tissue was chopped into small pieces in RPMI with 25 mM HEPES and 2% FCS. Following centrifugation, supernatant was removed and cells digested in collagenase solution (0.5 mg/mL collagenase type D, 0.2 mg/mL DNase I and 2% FCS in RPMI) at room temperature for 30 min with constant shaking. Digested tissue was filtered and then washed in 2% FCS.

#### Skin

dorsal and ventral half of each ear were separated, minced and left to incubate in collagenase solution (3mg/mL collagenase IV, 0.1 mg/mL DNAse I, 0.5 mg/mL hyaluronidase) for 30 min at 37°C with constant shaking. Digested tissue was filtered and washed in 2% FCS.

#### Tumors

tumors were excised, with skin and fat carefully removed. They were minced in collagenase solution (1 mg/mL collagenase IV, 20 μg/mL DNAse I and 2% FCS) and incubated at 37°C with constant shaking for 20 min. Digested tissues were filtered and washed in 2% FCS.

#### Synovial fluid

synovial fluid was obtained by puncturing the inflamed synovium with a 25G needle and collecting the extracted fluid into DMEM with 5% FBS and 20 mM EDTA.

### Flow cytometry

Cells were stained with the following surface marker antibodies: anti-mouse CD45 (BV510™, BioLegend, clone 30-F11, cat# 103138), anti-mouse CD11c (BV785™, BioLegend, clone N418, cat#117336), anti-mouse/human CD11b (Pacific Blue™, BioLegend, clone M1170, cat# 101224), anti-mouse F4/80 (APC-Cyanine® 7, BioLegend, clone BM8, cat#123118), anti-mouse TCRβ (BUV737®, BD Bioscience, clone H57-597, cat# 612821), TCRγδ (PE-Cyanine® 7, BioLegend, clone GL3, cat# 118124), anti-mouse CD4 (BV711™, BioLegend, clone RM4-5, cat# 100548), anti-mouse CD8a (AlexaFluor® 700, BioLegend, clone 53-6.7, cat# 100730), anti-mouse CD25 (APC, BioLegend, clone PC61, cat# 102012), anti-mouse CD19 (BV605™, BioLegend, clone 6D5, cat# 115540). Cells were stained for 25 min at 4°C, washed, and then acquired on the BD FACSymphony. Analysis was performed with FlowJo software. Flow t-SNEs were generated with the t-SNE FlowJo package. Chord diagrams of flow quantifications were generated using the Circlize package in R ^68^

### Single cell RNA-seq

Single cell RNAseq experiments were performed and analysed as previously described ^37^. Cecum cell suspensions were stained on ice for 20 min with anti-CD45, CD4, CD19, CD8, CD11b, CD11c, γδTCR, NK1.1 and DAPI as a viability dye. Cells were sorted as CD45^+^ DAPI^-^. Additional sorting was performed for CD19^-^ CD45^+^, NK1.1^+^ CD45^+^, and TCRγδ^+^ CD45^+^ to enrich for these populations. Splenic cell suspensions were stained with anti-CD45 (BioLegend) and DAPI as a viability dye (BioLegend). Kaede green and Kaede red cells from each mouse were sorted as CD45^+^ DAPI^-^, into 0.04% BSA. TotalSeq-A hashtag antibodies (0.5μL/sample for 20 min, BioLegend; hashtag 1, 24hr Kaede green mouse #1; hashtag 2, 24hr Kaede green mouse #2; hashtag 3, 48hr Kaede green mouse #3; hashtag 4, 48hr Kaede green mouse #4; hashtag 5, 72hr Kaede green mouse #5; hashtag 6, 72hr Kaede green mouse #6; hashtag 7, 24hr Kaede red mouse #1; hashtag 8, 24hr Kaede red mouse #2; hashtag 9, 48hr Kaede red mouse #3; hashtag 10, 48hr Kaede red mouse #4; hashtag 11, 72hr Kaede red mouse #5; hashtag 12, 72hr Kaede red mouse #6) were added to each sample after sorting. All samples were washed and then pooled together, centrifuged, and resuspended in 0.04% BSA. Encapsulation was done on the 10X Chromium microfluidic instrument (10X Genomics). Libraries were prepared using Chromium Single cell 3’ reagents kit v2 according to manufacturer’s protocol. Hashtag oligonucleotide (HTO) libraries were prepared as described in ^36^. Libraries were sequenced together on the Illumina HiSeq X. Cecum single cell was performed independently from the spleen.

#### Data analysis

scRNAseq data were processed using the standard CellRanger pipeline (10X Genomics). HTO counts were obtained using the CITE-seq-Count package ^69^. Data was analysed in R using the Seurat package ^70^. HTOs were assigned to cells using the HTODemux function, and doublets were eliminated from analysis. Cells with less than 700 UMIs or 500 genes and more than 2,500 UMIs, 10,000 genes and 5% of reads mapped to mitochondrial genes were also excluded from the analysis. Dimensionality reduction, visualization and clustering analysis were performed in Seurat using the NormalizeData, ScaleData, FindVariableGenes, RunPCA, FindNeighbours (dims=1:30), RunUMAP (dims=1:30) and FindClusters functions. Cluster identity was determined based on expression of key marker genes (Fig. S3A). The SubsetData function was used to remove individual B and T cell clusters for further analysis. Differentially expressed genes between Kaede red and Kaede green were obtained using the FindMarkers function on each of the top four Kaede red-enriched clusters and non-redundantly collated using a for-loop for each timepoint. The top 200 upregulated and downregulated genes across each of the three lists were selected and then also collated resulting in a final differentially list of genes generated based on statistical analysis (t-test, p<0.01). Heatmaps of differentially expressed genes were generated using Morpheus (https://software.broadinstitute.org/morpheus). Visualization of differential cell expression between Kaede red and Kaede green cells was done using the Buencolors package. Reference mapping of the Kaede red cells onto a colonic reference dataset was performed following Seurat’s data integration tool ^45^, using the FindTransferAnchors and MapQuery functions.

To identify iNKT cells expressing the canonical TCR, TCR V regions expressed in each cell were assembled using the TRUST4 algorithm ^71^ and searched for cells utilizing the canonical TRAV11/TRAJ18 combination; 27 such cells were found in the dataset, all of which shared the typical VVGDRGSAL CDR3a sequence, and all mapped to the NKT cluster of Fig.4A.

#### Gene signatures

The IEL gene signature was based on the expression of the following marker genes: *Klra1, Klre1, Klra7, Itgae, Cd160, Klrk1, Fasl, Itgb7, Ccr9, Cd8a*. The NKT gene signature was on the expression of the following marker genes: *Il4, Atp8a2, Trpm6, Garnl3, Hpn, Apt6v0d2, Klrb1c, B3galt5, Gpnmb, Rnf144b, Serpina3f, Zbtb16, Klra1, Sulf2, Rin2, Art2a-ps, Arg1, Gm9195, Pls1, Wnt10a, Fam84a, Csf2, Zfp683, Mgll, Apol7b, Dgki, Cd93, Fbxl21, Fcrls, Chad, Flt4, Rmdn2, Klrb1b, Ckb*.

## Statistical analysis

Statistical analyses outside of single cell data were performed using GraphPad Prism software. Unless stated otherwise, data is presented as mean ± SD. Statistical significance was calculated by unpaired Student’s t-test. P < 0.05 was considered significant.

## Data and materials availability

Data newly reported in this paper were deposited in the Gene Expression Omnibus (GEO) database under accession number xxx.

## SUPPLEMENTARY FIGURE LEGENDS

**Figure S1.**
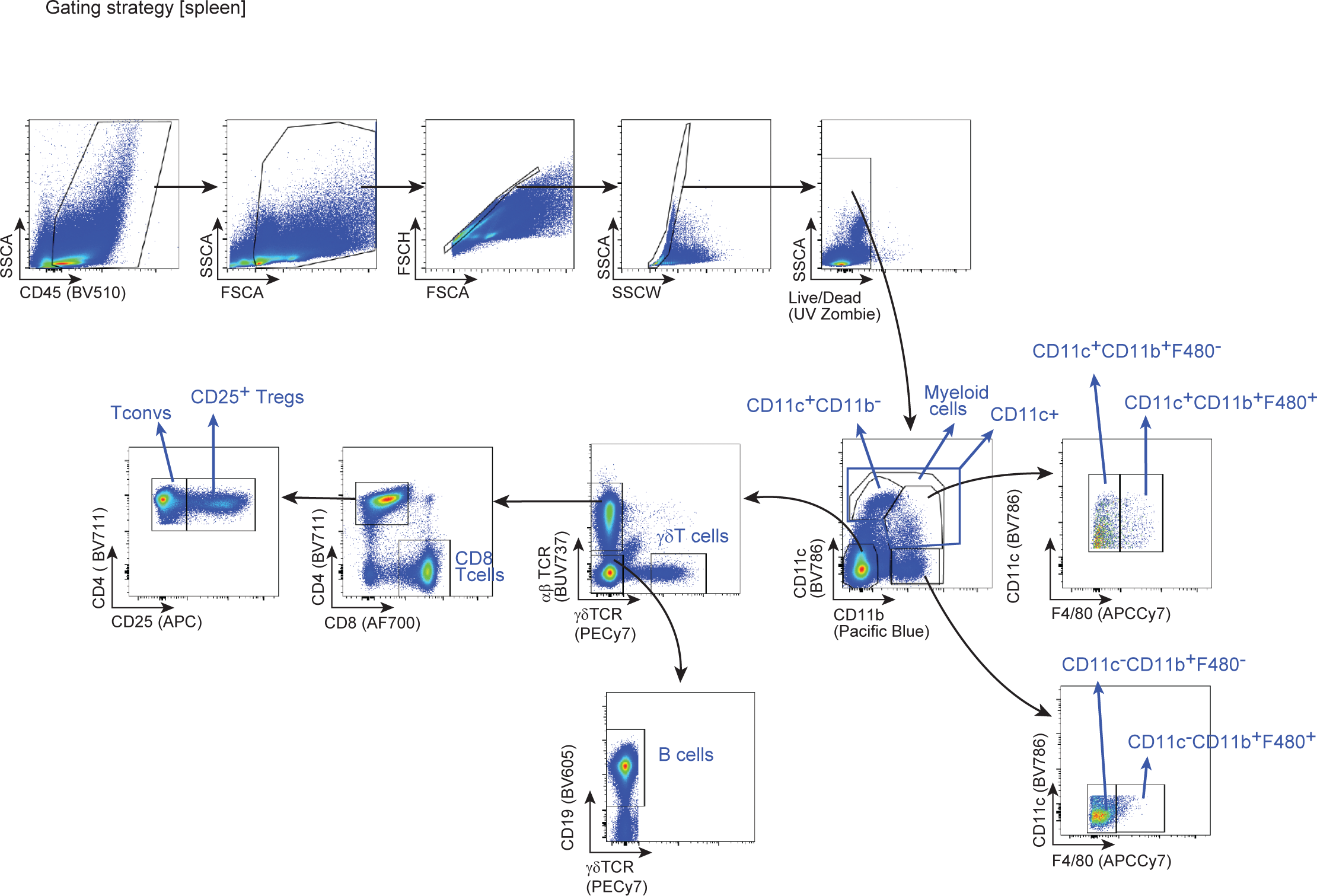
Flow cytometry gating strategy. Representative flow cytometry plots of the gating strategy for the studied immunocyte populations.

**Figure S2.**
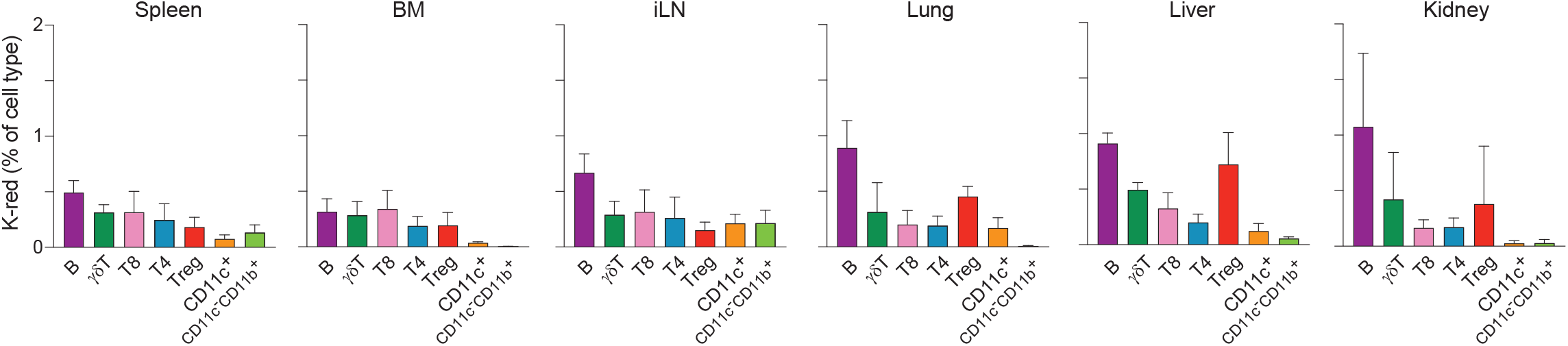
Representation of colon-derived immunocytes across multiple tissues. Percentage of immunocyte populations in peripheral tissues 24hr post-photoconversion of the descending colon, as measured by flow cytometry.

**Figure S3.**
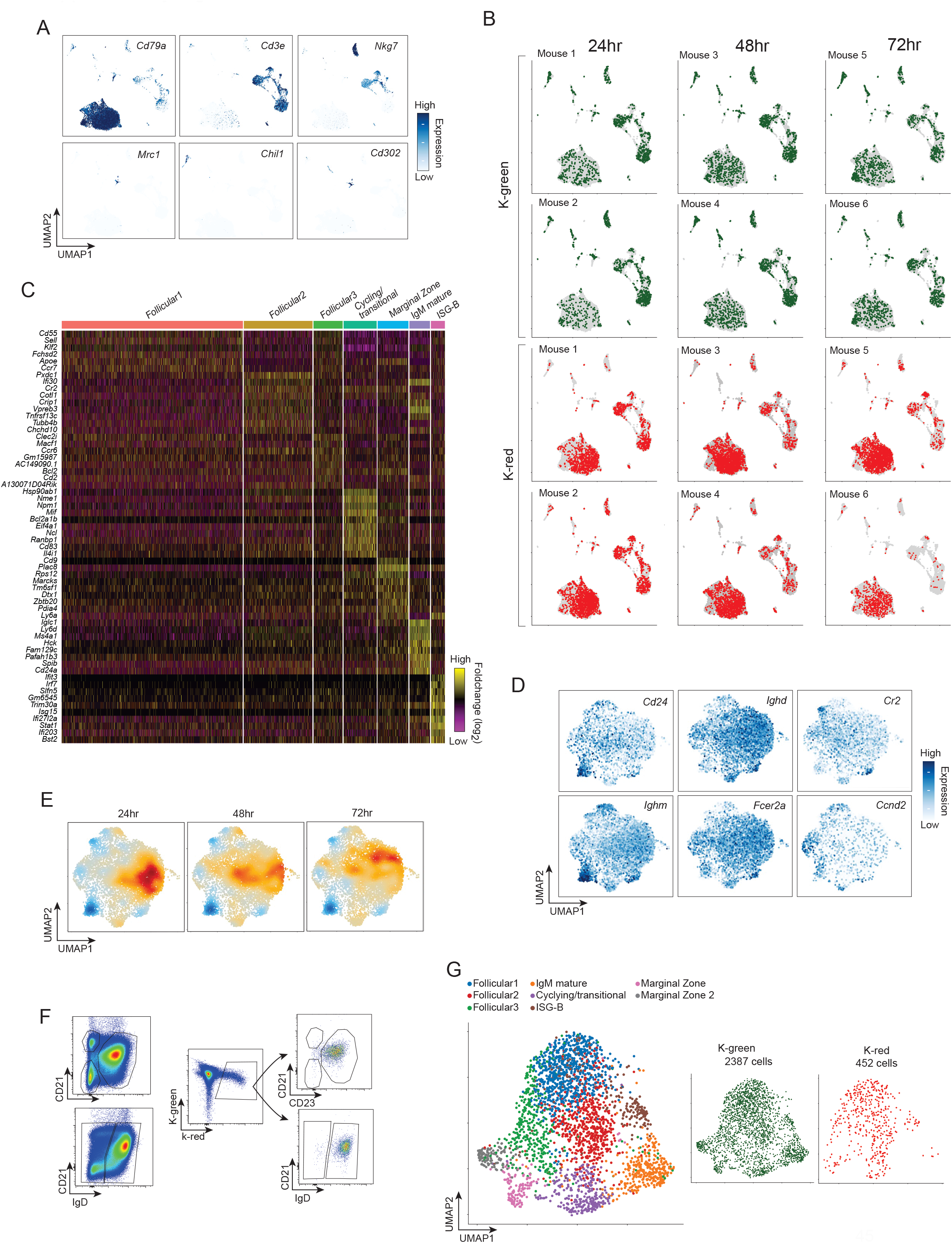
scRNAseq analysis of local and colon-derived CD45^+^ cells in spleen. A) Same UMAP as in Fig. 3B, with gene expression of selected markers highlighted. B) Same UMAP as in Fig. 3B, with cell identity highlighted as determined by mouse, timepoint and Kaede-green vs Kaede-red. C) Heatmap of most differentially expressed genes across each B cell cluster, depicted in Fig. 3C. D) Same UMAP as in Fig. 3C, with gene expression of selected markers highlighted. E) Same UMAP as in Fig. 3C, with a differential density representation of Kaede-red (red) vs Kaede-green cells (blue) across each timepoint. F) Confirmation of marker expression by flow cytometry (gated on Live, CD45^+^ CD3^-^ CD11b^-^ CD11c^-^ CD19^+^ cells). G) Single cell RNAseq analysis of splenic B cells using the same set-up but performed independently from the one in Fig. 3. Left panel depicts the identified B cell clusters, right panels depict the distribution of the Kaede green vs Kaede red across the same UMAP.

**Figure S4.**
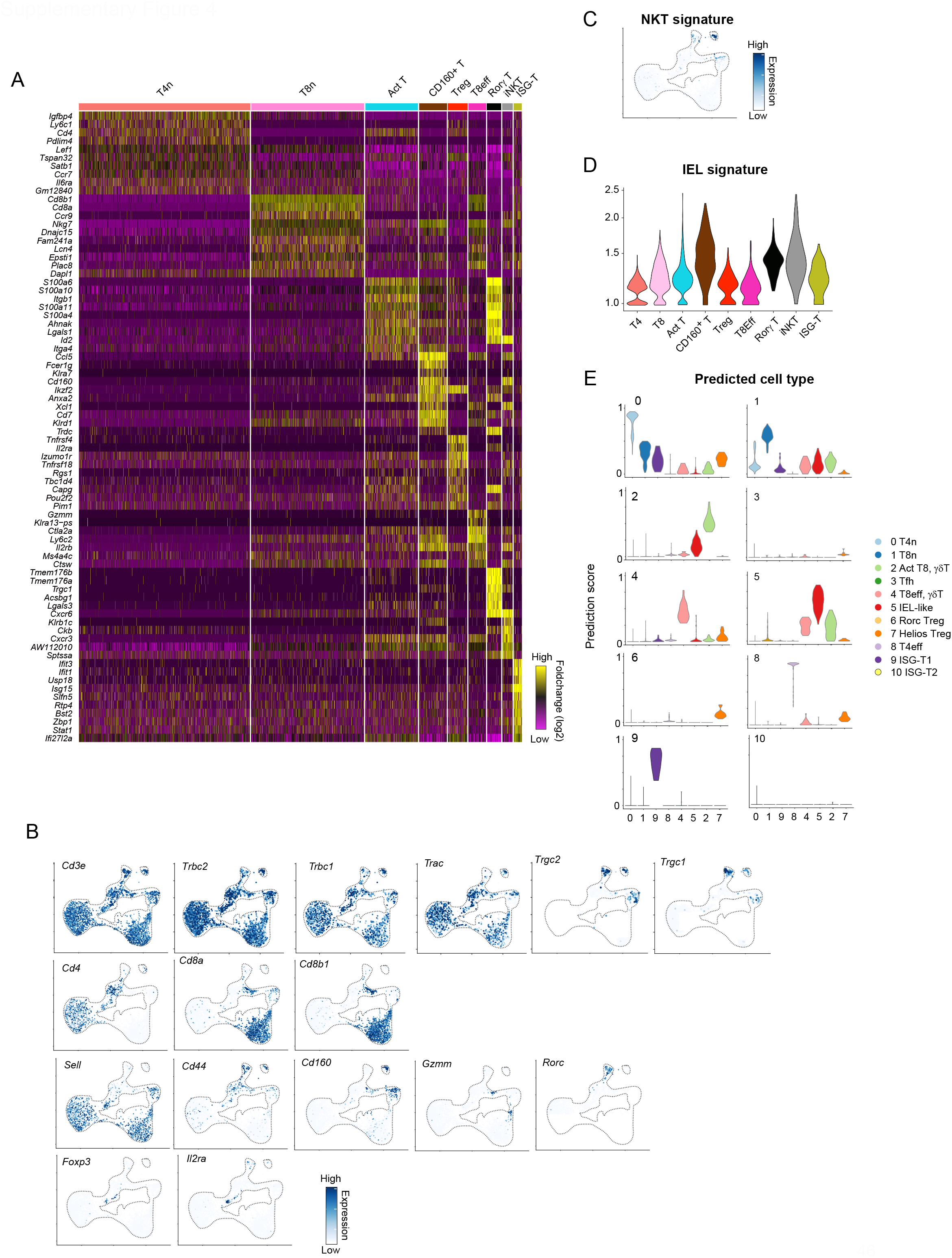
scRNAseq analysis of local and colon-derived T cells in spleen. A) Heatmap of most differentially expressed genes across each T cell cluster, depicted in Fig. 4A. B, C) Same UMAP as in Fig. 4A, with gene expression of selected markers (B) and a gene signature of NKT cells (C) highlighted. D) Expression of an IEL gene signature across T cell clusters from 4A. E) Prediction scores used to for cluster assignment of cells in Fig. 4G.

**Figure S5.**
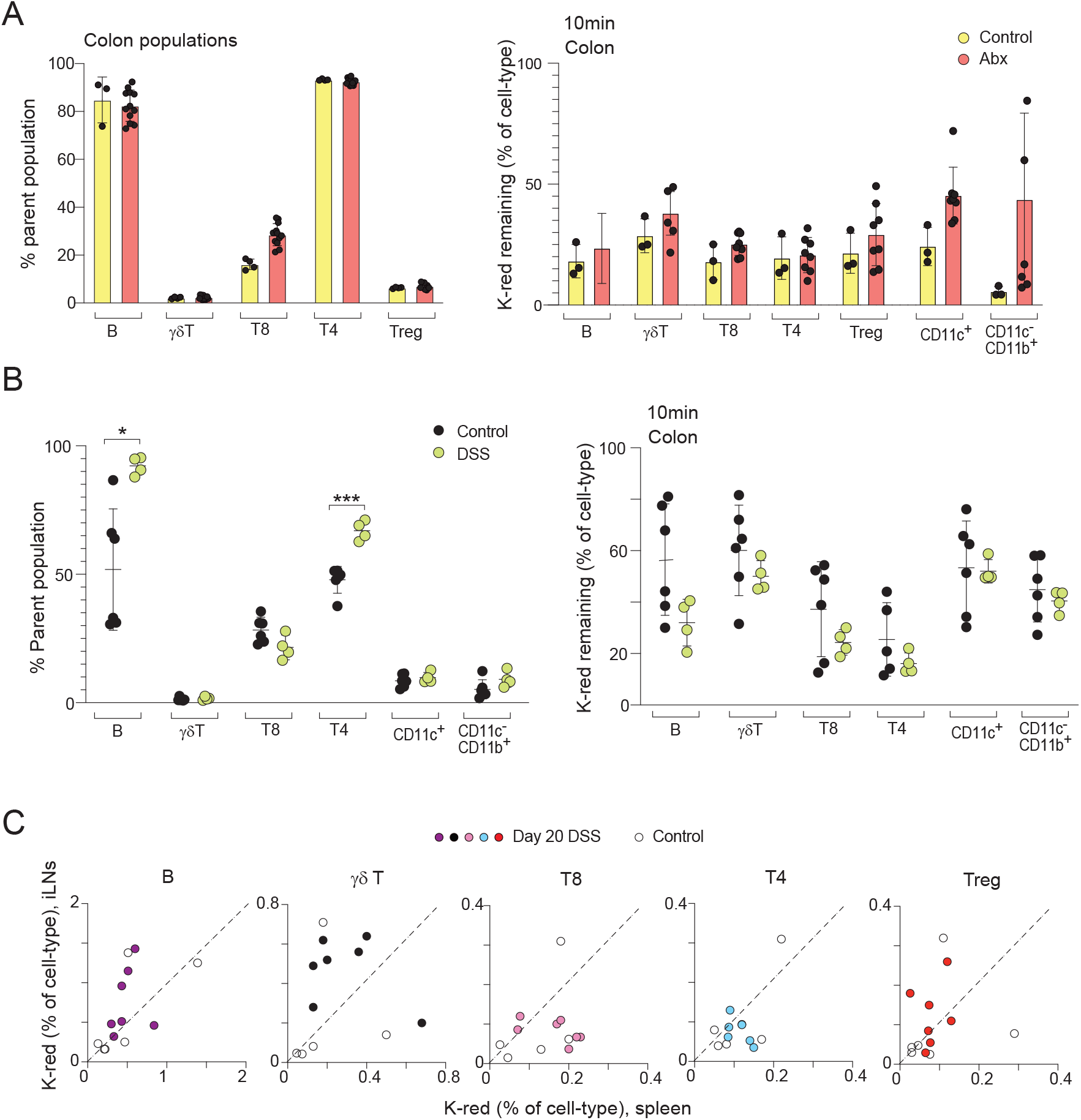
Immunocyte population dynamics and photoconversion in antibiotic or DSS-treated mice. A) Proportions of immunocyte populations (left) and photo-tagged immunocyte populations remaining in the colon (right), 10 min post-colonic photoconversion in control and antibiotic-treated mice (10 days, VGCA). B) Proportions of immunocyte populations (left) and photo-tagged immunocyte populations remaining in the colon (right), 10 min post-colonic photoconversion in control and DSS-treated mice (7 days). C) Correlation between Kaede-red immunocyte populations in iLN (y-axis) and in spleen (x-axis) in control vs mice on day 20 post initial DSS administration, 24hr post-photoconversion of the descending colon. All results from two to four independent experiments. Each dot represents a mouse, mean is marked. *p<0.05; ***p<0.001, unpaired t-test.

**Figure S6.**
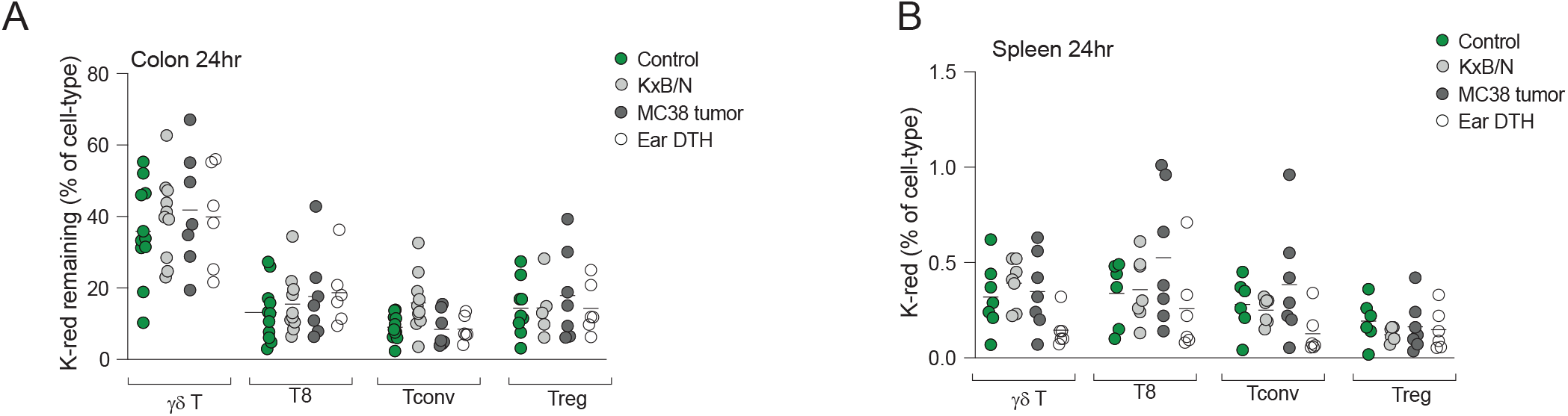
Colon egress and spleen migration of T cell populations in arthritis, tumor and skin hypersensitivity. A) Proportions of T cell populations left in the descending colon 24hr post-photoconversion in control and diseased mice. B) Percentage of colon-derived T cell populations in the spleen 24hr post-photoconversion of the descending colon in control and diseased mice. All results from two to four independent experiments. Each dot represents a mouse, mean is marked.

